# Analysis of the spatiotemporal dynamics of behavior in domestic dogs with free locomotion under appetitive Pavlovian contingencies

**DOI:** 10.64898/2026.02.20.707116

**Authors:** Abraham Rivera, Varsovia Hernández, Danna Jiménez, Alejandro León

## Abstract

Spatiotemporal dynamics of behavior is key to understanding the organism-environment relationship. While often implicitly addressed, its relation to Pavlovian contingencies remains understudied in domestic dogs. This study quantitatively examined the spatial dimension of behavior of three freely moving dogs under pairing and extinction of a tone–food Pavlovian contingency. In pairing, a tone (CS) was paired with food delivery (US) on a fixed-time 60-second (FT 60 s) schedule; in extinction, only the tone was presented. Locomotion was recorded using two-dimensional tracking based on center of mass. During pairing, dogs moved closer to the dispenser, covered greater distances, displayed extended trajectories, and showed a conditional approach pattern to the dispenser during CS presentation. In extinction, they stayed closer to the owner or room periphery, traveled shorter distances, and exhibited more restricted trajectories. These findings show that spatial segments (dispenser area) are integrated into the CS–US relationship, demonstrate the usefulness of continuous recording of spatial behavior in the analysis of Pavlovian contingencies, and suggest potential application in contexts relevant to animal welfare.

**Highlights:**

- Spatial behavior remains largely unexplored in studies of Pavlovian contingencies
- Dog locomotion was continuously tracked during pairing and extinction phases
- Distinct spatial patterns emerged across pairing and extinction phases
- Incorporating space into CS–US relations clarifies behavioral organization
- Tracking tools expand the experimental and applied scope of Pavlovian research

## 1 Introduction

The spatiotemporal dynamics of behavior have been identified as a crucial dimension in understanding the organism–environment relationship (León et al., 2021). This dimension is observed in the emerging patterns of an organism’s continuous movement in relation to the occurrence of specific events, as well as in the structural arrangement of the environment in which the behavior occurs.

The study of the spatiotemporal dynamics of behavior is fundamental to understanding how behavioral patterns emerge in organisms exposed to contingency arrangements that depend on temporal criteria, particularly during intervals when discrete responses do not occur (Pear, 1985). These arrangements, which integrate temporal criteria for the presentation of events (e.g., delivery of reinforcers or environmental changes), include both reinforcement schedules that combine the passage of time with the occurrence of responses (e.g., Interval Schedules) and those that are governed exclusively by time (e.g., Time Schedules). Hereinafter, both will be referred to as temporal arrangements.

Various studies have shown that modifications in temporal arrangements can give rise to distinctive spatial behavior patterns in several species (Pear, 1985; León et al., 2020, 2021). For example, in pigeons, displacement patterns tend to change depending on modifications in the temporal parameters of Variable Interval (VI) programs. In 5-minute VI programs, extended movement trajectories and patterns of going back and forth to the operant have been reported, while in 15-s VI programs, movement patterns tend to be more confined to areas close to the operant (Pear, 1985).

Likewise, studies with rats (León et al., 2020) have reported quantitative variations in movement patterns depending on different types of temporal programs, for example, Fixed Time (FT) and Variable Time (VT). In particular, it has been observed that VT programs tend to generate a more pronounced decrease in spatial dynamics compared to FT programs. These findings underscore the importance of analyzing the interaction between the temporal and spatial dimensions of behavior, providing evidence on how organisms adjust the dynamics of their movement based on the spatiotemporal conditions of their environment.

Pavlovian contingencies — also called classical conditioning procedures — are some of the most studied temporal arrangements (Domjan, 2010). These procedures involve the con-ditional rela- tionship between the presence of an environmental stimulus (neutral stimulus that becomes a conditioned stimulus, CS) and the subsequent appearance of an unconditioned reinforcing stimulus (unconditioned stimulus, US) that reliably elicits a response (unconditioned response, UR). The frequency, duration, and latency of responses elicited by the CS are the main dependent variables when studying this type of contingency arrangement (Henton and Iversen, 1978).

Although the relationship between Pavlovian contingencies and the spatiotemporal dimension of behavior is not often emphasized, probably because the initial preparation involved working with immobilized subjects (Pavlov, 1927), this dimension has been implicitly explored in various studies over the years (Kupalov, 1969; Kupalov et al., 1964; Kolosova et al., 1982, among others).

The literature pertaining to the areas of autoshaping (Brown and Jenkins, 1968), sign-tracking behavior (Hearst, 1975; Jenkins et al., 1978), and place conditioned or situational conditioned reflexes (Kupalov, 1969; Kupalov et al., 1964; Kolosova et al., 1982) exemplify the historical interest in studying spatiotemporal dynamics in freely moving organisms exposed to Pavlovian contingencies. This research not only highlights the wide range of behaviors that can be part of a functional organism–environment relationship, but also provides empirical evidence to support the assumption that a particular behavioral organization (established from a specific contingency arrangement) does not occur in a spatial vacuum, but rather that space itself is a fundamental element that will participate in that relationship (Kupalov, 1969, 1983).

Kupalov (1969) was one of the first researchers to conduct a formal analysis of the spatial behavior of organisms by studying the formation of situational conditional reflexes (Buzsáki, 1982). In his analyses, he considered locomotion patterns and various behavioral morphologies in dogs (e.g., posture, location, orientation, and movement of the organism). The experimental setup used by Kupalov (1969) consisted of a large room containing two separate tables, each with a food dispenser and various types of CS (bell, buzzer, lights, tones, and sounds of varying frequencies and intensities). The procedure began with an initial CS-US pairing (e.g., food) at each table. Once the subjects consistently moved to each table when the CS was presented, the CS was presented contingently on the dog’s location within a specific area of the experimental space (indicated by a mat).

The results of Kupalov’s study (1969) showed the narrowing of movement patterns between the mat area and the tables, as well as the emergence of particular behaviors (e.g., postures or orientations) over the course of the sessions, such that the dogs’ behavior became increasingly consistent and with fewer movements irrelevant to obtaining the US. For Kupalov (1969; 1983), the interaction of spatial variables and stimulus–stimulus contingency relationships formed a situational conditional reflex. When the place or situational conditional reflex was firmly established, a specific place in space became a positive conditional agent, so that the animal actively and independently went to this place, adopting a specific posture and remaining there until the subsequent CS was presented.

Although Kupalov’s findings (Kupalov, 1969, 1983; Kupalov et al., 1964) were of great importance due to their methodological innovation and consideration of the spatiotemporal dimension of behavior in the study of conditioned reflexes, it has been pointed out that his methodological paradigm is basically an operant paradigm (Buzsáki, 1982), because the presentation of the trials (CS-US) ultimately depended on the location of the dogs on the mat. Because of this, it is difficult to attribute the establishment of behavioral organization to the effect of Pavlovian contingencies per se rather than to interaction with operant processes.

This discretization of trials, based on the fulfillment of a criterion of the dogs’ own activity, remained a recurring procedure in the study of situational and place conditioned reflexes (e.g., Kolosova et al., 1982). Even in research on sign-tracking and goal-tracking with dogs (Jenkins et al., 1978; Stepień, 1974), a subtle discretization of trials based on the animal’s position on a starting mat has been used, which continues to make it difficult to distinguish the effects of Pavlovian contingencies from operant contingencies.

Additionally, it is important to note that, unlike previous studies with other species (e.g., Hearst et al., 1980; León et al., 2020; Pear, 1985; Wasserman et al., 1974), quantitative analysis of spatiotemporal dynamics in dogs under Pavlovian contingencies has been poorly explored. Analyses of spatiotemporal dynamics in these studies have been based on observations by experimenters, behavioral descriptions, and pictorial representations of movement patterns (e.g., Jenkins et al., 1978; Kupalov, 1969; Stepień, 1974; Zener, 1937).

In this regard, most studies with dogs under Pavlovian contingencies have not accurately, automatically, and quantifiably recorded the movement of dogs in the experimental space, which is now achievable using contemporary computational technologies for tracking deformable objects (e.g., image processing-based tracking systems) that have already been used in other fields of research with the same species (Barnard et al., 2016; Bleuer-Elsner et al., 2019; Völter et al., 2023; Zamansky et al., 2018). These tools would allow for the incorporation of variables that are not usually analyzed, such as the intensity, distribution, and variability of behavior in space in relation to the occurrence of relevant events (e.g., León et al., 2020, 2021).

Likewise, this type of analysis would allow us to transcend the analytical limitations of an approach focused exclusively on the organism’s proximity and contact with the sources of stimulation established by the experimenters (as in the case of research on sign-tracking or goal-tracking), since it makes it possible to measure behavior independently of a reference point (such as movement routes, total distance traveled, location entropy, among others).

Based on the above, this study analyzes the spatiotemporal dynamics of behavior in domestic dogs under appetitive Pavlovian contingencies, performing a quantitative assessment of behavior (e.g., movement) in space using tracking tools, as well as robust spatial data analysis (León et al., 2021). The hypothesis is that, under Pavlovian appetitive contingencies, the regions of space associated with the dispenser may acquire a conditional function (Kupalov, 1969, 1983), organizing the spatiotemporal dynamics of behavior.

Unlike previous studies in which the choice of domestic dogs as experimental subjects is not explicit (e.g., Jenkins et al., 1978; Kupalov, 1969), in the present study the choice was based on: 1) the ease of studying basic behavioral processes in this species; 2) future comparison among species; and 3) the possibility of extending results to applied scenarios that seek to improve the quality of life of domestic dogs (Fernandez et al., 2023; Feuerbacher and Wynne, 2011; Lindsay, 2000; Protopopova and Wynne, 2015).

## 2 Method

### 2.1 Experimental subjects

The experimental subjects were three experimentally naive domestic dogs: two mixed breeds (one ∼6-year-old male and a ∼4-year-old female) and a pit bull (∼7-year-old male). The animals were deprived of food for ∼12 hours (12 ≤ *x* < 24 hours). The deprivation period was adjusted to the dogs’ usual feeding schedule, so no changes were made to the dogs’ feeding routine. At the end of each experimental session, each dog was fed normally by its respective owner at its own home.

The selected dogs were domestic pets that met the inclusion criteria. They were experimentally naive dogs with no previous experience with the stimuli used; they lived mainly indoors in close contact with their owners; they were clinically healthy, with no apparent diseases or conditions that affected their interaction with their environment (e.g., blindness, deafness, or mutilation). All dogs were within an age range of two to eight years and showed a willingness to consume the kibble used. Additionally, the selected dogs were already on a ∼12-hour food deprivation regimen imposed by their owners.

The owners signed an informed consent form for holders of research animals. In addition, prior to the start of the experiment, they completed a questionnaire about the animal’s welfare. The questionnaire served as an additional filter that allowed researchers to identify signs of undiagnosed diseases while the experiment was being conducted, thereby reducing the likelihood of including dogs with health problems.

All procedures were approved by the Internal Committee for the Care and Use of Laboratory Animals (Approval No. CLCIB2024/2) and were carried out in accordance with Mexican standard NOM-062-ZOO-1999 for the production, care, and use of laboratory animals. Although this study did not involve laboratory animals, the ethical principles and guidelines established in this standard were followed to ensure the welfare of the experimental subjects in this research.

### 2.2 Equipment and experimental setup

The study was conducted in a domestic-naturalistic setting. The decision to conduct this study in domestic-naturalistic settings, i.e., in rooms shared with their owners, is supported by recent studies that have made behavioral observations in natural situations and away from traditional experimental measures, such as behavioral observations in parks (e.g., Howse et al., 2018) and reinforcement procedures in homes (e.g., Mehrkam et al., 2020). This experimental methodology not only offers greater ecological validity, but also allows research to be conducted in an ethical context, where manipulation and intervention by the experimenter is minimized, thus prioritizing animal welfare.

Based on this methodology, a room measuring 2.80 m × 2.66 m was adapted to carry out the experimental sessions. A chair was placed inside the room so that the owner could sit down during each experimental session. The owner remained inside the room during all experimental sessions in order to minimize the adverse effects of animal isolation.

An automatic food dispenser (30 × 28 × 30 cm) with a built-in buzzer was used to deliver food and play a sound tone. The dispenser consisted of an Arduino^®^ Uno microcontroller, an actuator (PAP Unipolar 28BYJ-18 5V motor, ULN2003 driver module) that operated a perforated MDF disc, and a PVC delivery chute. The tones used as a conditional stimulus (23 kHz, 75 dB) were emitted from an active buzzer module adapted to the dispenser.

The food used as an unconditional stimulus for the pairing sessions was Iron Dog^®^ premium dog food. This food was chosen based on its nutritional standards and its suitability for the dispenser. Each delivery consisted of the release of one kibble that fell onto the floor. A cloth patch (30 cm × 30 cm) was used, which was positioned just below the dispenser’s delivery chute to reduce the sound produced and the movement of the kibble as it fell.

During the experimental sessions, the experimenter remained outside the room, operating the recording and control of experimental events using software developed in Visual Studio^®^ 2015.

The location of the dispenser (X, Y) and the owner (Z, W) remained fixed in all sessions. All sessions were recorded on video with an RGB camera (1920 × 1080, 10 fps) placed on the ceiling of the room, which remained in the same location throughout the experiment. The layout of the experimental setup can be seen in Figure 1.

**Figure 1.**
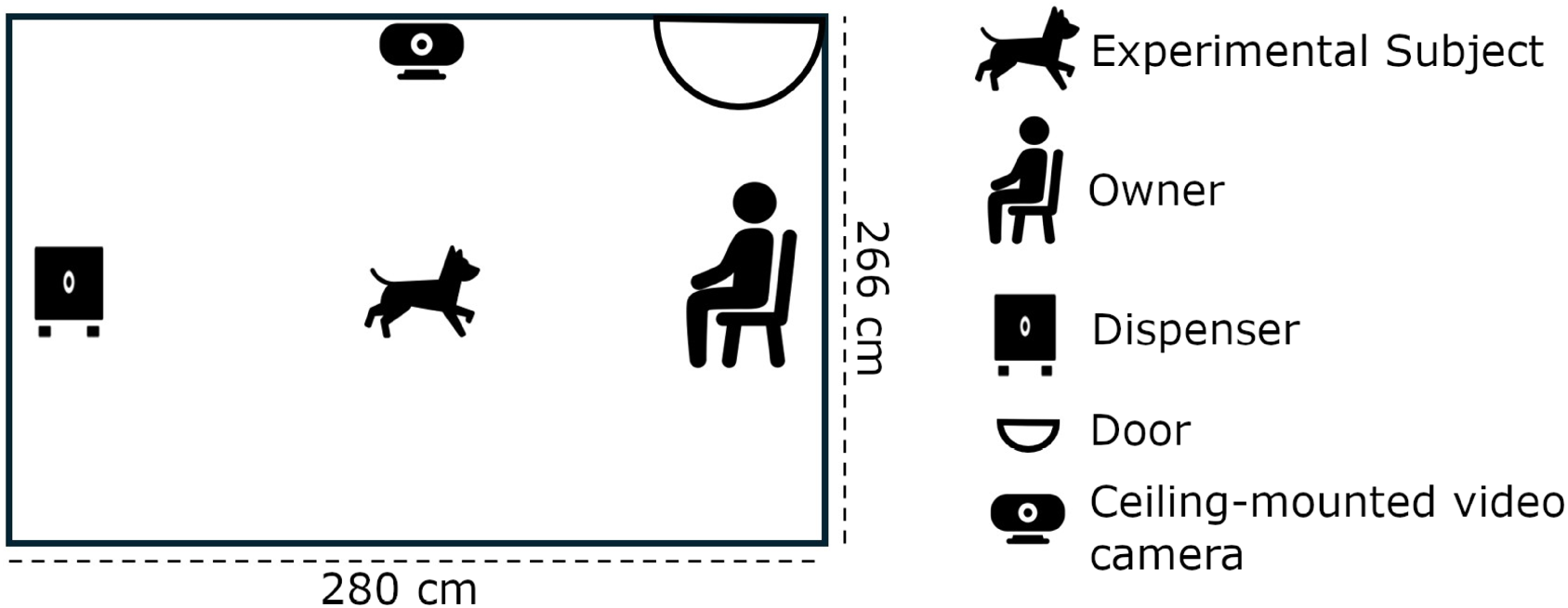
Experimental setup.

### 2.3 Design

A single-case reversal (B–A–B–A) design, replicated across three subjects, was used. In condition B (Pairing), a tone was paired with the delivery of food. During condition A (Extinction), only the tone was presented without the delivery of food. The experimental design is outlined in Table 1. The extinction phase was included because it allowed for a comparison of the behavioral changes generated by the alteration of the contingency relationship with those generated by the CS-US pairing, as extinction phases are typically incorporated in Pavlovian conditioning procedures (Tarpy, 2000).

**Table 1:**
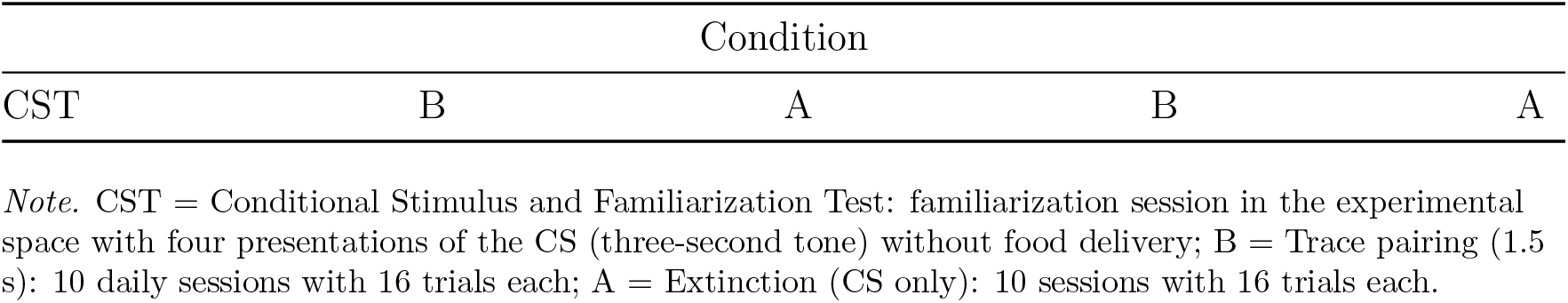
Experimental design Condition.

| Condition |  |  |  |  |
| --- | --- | --- | --- | --- |
| CST | B | A | B | A |

*Note.* CST = Conditional Stimulus and Familiarization Test: familiarization session in the experimental space with four presentations of the CS (three-second tone) without food delivery; B = Trace pairing (1.5 s): 10 daily sessions with 16 trials each; A = Extinction (CS only): 10 sessions with 16 trials each.

### 2.4 Procedure

The experimental sessions were held Monday through Friday, between 6:00 p.m. and 9:00 p.m., for all subjects. Each session was conducted individually, and at no time did the dogs or their respective owners have contact with other participants.

In each session, the owner brought their dog to the experimental room, where both remained. The dog moved freely within the space, while the owner remained in the designated area. To minimize the emission of inadvertent signals, the owner was instructed not to speak, make noises, or make eye contact with the animal throughout the session. The dogs were exposed to two types of conditions: Pairing (B) and Extinction (A).

Before the experiment began, the subjects underwent a familiarization session lasting 16 minutes and 55 seconds. During this session, each dog remained in the experimental room, while its owner stayed in the designated area. The subjects were free to move around. In the last four minutes of the session, a three-second tone (23 kHz, 75 dB) was automatically emitted through a dispenser located in the room, repeating four times every 60 seconds. The tone was the same as that used later in the pairing and extinction conditions. This brief exposure to the auditory stimulus was intended to identify orientation responses in the subjects evoked by the tone prior to pairing. Its implementation made it possible to distinguish the behavioral patterns promoted by the CS–US correlation from those promoted solely by the tone and the experimental arrangement. Hereafter, this initial session will be referred to by the acronym CST (Conditional Stimulus and Familiarization Test).

#### *Pairing* (B)

During pairing, the automatic delivery of a treat was programmed on a fixed 60-second timer (FT60 s). In each trial, the dispenser emitted a three-second tone, and one and a half seconds after it ended, the treat was released (trace conditioning). This condition consisted of 10 sessions, each session consisting of 16 consecutive trials. The duration of each session was 16 minutes and 55 seconds.

#### *Extinction* (A)

The extinction condition consisted of 10 sessions. The parameters were identical to those of the pairing condition, except that the tone was not followed by food delivery.

Once the Pairing–Extinction (BA) sequence was completed, they were replicated in the same order and in the same way as described above (BABA). To facilitate identification and tracking, each dog was assigned a distinctive-colored garment worn during all experimental sessions.

### 2.5 Data recording and analysis

The video recordings of the experimental sessions were processed using Ethovision XT software (version 14), which was used to obtain the coordinates (X, Y) of the center of mass of the experimental subjects at a resolution of 10 Hz.

For data analysis, Motus and R (version 4.3.3) programs were used within the RStudio environment. The variables analyzed included movement paths, cumulative time (frames) in regions, total distance traveled, distance to relevant objects (dispenser-owner), conditional approach index to the dispenser (Hearst et al., 1980; Wasserman et al., 1974), and location entropy (León et al., 2021).

The visualizations and measures used are described below:

#### Movement routes

The visualization of the movement routes was obtained by recording the X and Y coordinates of the subject’s center of mass and connecting them sequentially throughout the session, at a temporal resolution of 10 Hz, thus forming a continuous trajectory.

#### Time (frames) accumulated in regions

To calculate the cumulative time spent in different regions of the experimental space, the two-dimensional environment was divided into a virtual grid of 72 cells (9 columns × 8 rows). The X and Y coordinates of the organism that coincided with each of the resulting regions throughout the sessions (10 Hz resolution) were then identified.

#### Total distance traveled

The total distance traveled was calculated based on the Euclidean distance between consecutive coordinates. These values were then added together to obtain the total distance per session. The formula used was:

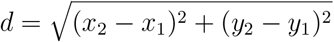

Where (*x*_1_, *y*_1_) and (*x*_2_, *y*_2_) correspond to the successive coordinates of the dog in space, and *d* represents the distance traveled between those coordinates.

#### Distance to relevant objects (dispenser-owner)

To obtain the distance values to relevant objects, the Euclidean distance between the dog’s location in the experimental room and the coordinates corresponding to the objects of interest (dispenser-owner) was calculated. These values were then averaged per session. The formula used was:

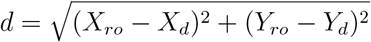

Where (*X*_*ro*_, *Y*_*ro*_) represent the coordinates of the relevant object (dispenser or owner), which remained constant throughout the experiment, while (*X*_*d*_, *Y*_*d*_) correspond to the coordinates of the dog. The value *d* represents the distance between the location of the dog and the location of the relevant object in each time unit, at a temporal resolution of 10 Hz.

#### Conditional approach index to the dispenser

A procedure based on previous research with pigeons (Hearst et al., 1980; Wasserman et al., 1974) was adopted. To this end, the experimental space was virtually divided into two mutually exclusive regions, segmenting the horizontal axis of the room exactly in half (140 cm). The area near the dispenser ranged from > 0 to 140 cm and included the location of the dispenser within its boundaries. On the other hand, the area far from the dispenser comprised the remaining region (> 140–280 cm), including the location of the owner. This segmentation allowed us to quantify the cumulative time of the subjects in each of the areas during the experimental sessions.

The conditional approach index to the dispenser (CAI) in each trial was defined as:

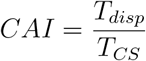

Where *T*_*disp*_ corresponds to the cumulative time spent in the area near the dispenser during the presentation of the CS, and *T*_*CS*_ corresponds to the total duration of the CS (30 frames). The index ranged from 0 (the dog was never in the area near the dispenser during the presentation of the CS) to 1 (the dog remained exclusively in that area for the entire duration of the CS).

#### Location entropy

Shannon entropy is a measure associated with discrete random variables and is useful for quantifying variability within a distribution (Carcassi et al., 2021; León et al., 2021). In the present study, the location entropy measure used was the same as that used in previous studies that have evaluated the spatial variability of rats in enlarged experimental arenas (León et al., 2020; Lanovaz et al., 2025).

According to Carcassi et al. (2021), entropy is formally defined as:

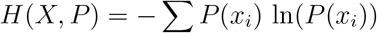

Where *X* is a discrete random variable with possible states {*x*_*i*_}, each with an associated probability *P* (*x*_*i*_).

In the context of this study, location entropy was used to evaluate the variability in the spatial distribution of the organism in each session (León et al., 2021). The discrete variable (*x*_*i*_) corresponded to the regions of the experimental space occupied by the organism, determined from its X and Y coordinates in each frame. These coordinates were assigned to one of the 72 regions defined by the virtual division of the two-dimensional space into a 9 × 8 grid. The probability *P* (*x*_*i*_) was obtained as the proportion of accumulated time (in frames) in each region. In this way, entropy quantified the distribution of the subject’s location, reaching higher values when the time spent was distributed evenly among several regions and lower values when the subject spent most of the time in a few areas (León et al., 2021).

Additionally, complementary statistical analyses were performed on the total distance traveled per session, location entropy per session, mean distance to the dispenser/owner per session, and conditional approach index to the dispenser per trial. For each variable, linear mixed models (LMM) were fitted, considering the experimental condition (pairing/extinction) as a fixed effect and the subject ID as a random intercept. The models were fitted in Jamovi (version 2.6.26) using the GAMLj3 module. In most cases, the residuals met the assumptions of normality, according to the Kolmogorov–Smirnov and Shapiro–Wilk tests (*p* >.05). Given that LMMs are robust to moderate deviations from this assumption (Schielzeth et al., 2020), the conditional approach index to the dispenser was retained in the analysis, even though it did not meet the assumptions.

## 3 Results

Figure 2 shows the movement paths of subjects (D1, D2, D3) in the CST, pairing (B1, B2), and extinction (A1, A2). For subjects D1 and D2, the paths correspond to session 10 of each exposure, while for D3, they correspond to the sixth session, since this subject showed low behavioral involvement in the subsequent pairing sessions, which were not considered representative. The brown squares and green triangles indicate the location of the dispenser and the owner, respectively. Additionally, the white–black color gradient represents the cumulative time in different regions of the experimental space. The regions with the highest cumulative time are represented with greater saturation. The graphs presented are an analog representation of the experimental room, where the solid black line shows the dog’s movement throughout the session.

**Figure 2.**
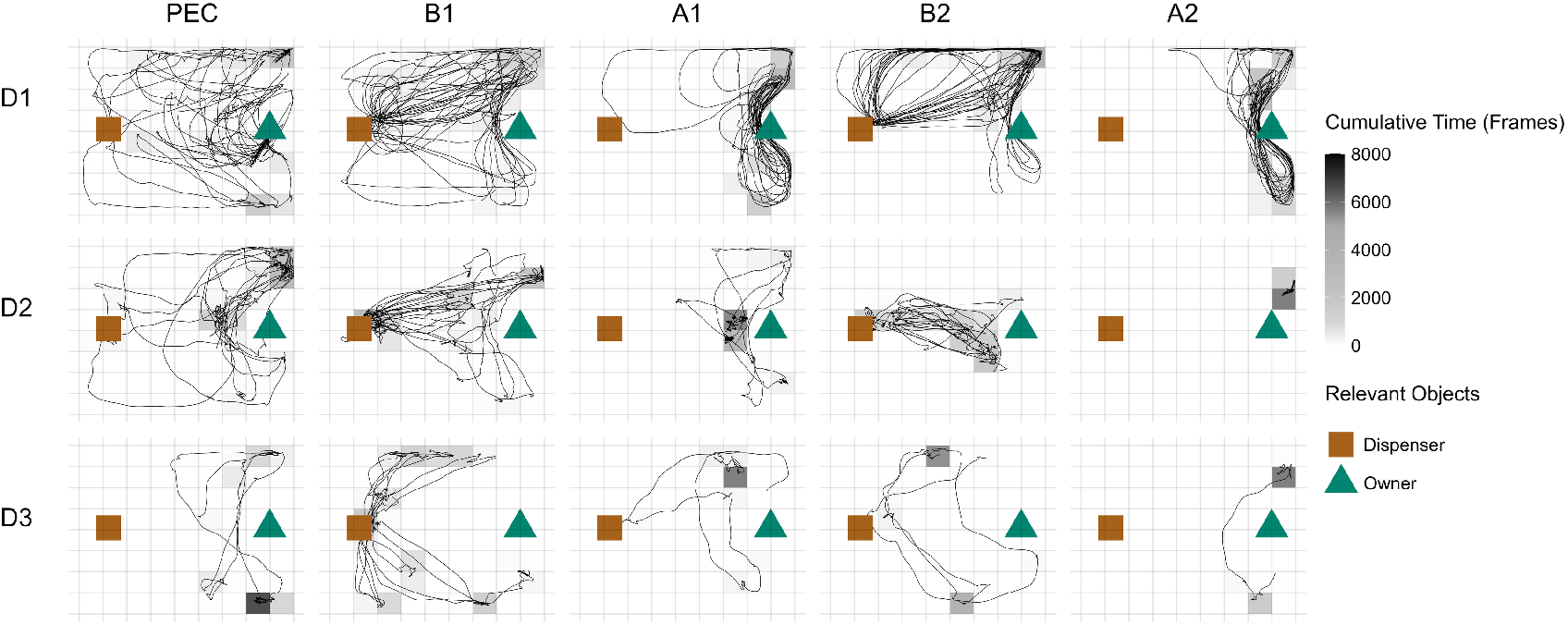
Movement paths during representative sessions. Each panel shows the analog paths recorded in representative sessions, where the brown square represents the location of the dispenser (XY coordinates: 28.5, 136) and the green triangle represents the location of the owner (XY coordinates: 248.5, 136). The color gradient, from white to black, indicates the cumulative number of frames in regions delimited by a 9 × 8 grid (0 to ≥8,000 frames). The rows correspond to each subject (D1, D2, D3), while the columns represent the experimental conditions—Conditional Stimulus Test and Familiarization (CST), Pairing (B) and Extinction (A)—ordered from left to right according to their sequence of exposure. The paths in B1, A1, B2, and A2 correspond to the last session of each exposure for D1 and D2, and to the sixth session for D3.

The owner was included as a relevant object due to the possibility that dogs would display activity patterns directed toward the owner, influenced by their history of interaction. Furthermore, it has been documented that the presence of the owner can increase levels of exploration in controlled contexts of free locomotion (e.g., Völter et al., 2023).

The routes in the CST became denser in areas close to the owner, while in the pairing condition (B1 and B2) a change in routes was observed in all dogs. In general, patterns of back-and-forth movement were established between areas close to the dispenser and peripheral areas of the room (e.g., upper right corner and walls) or regions close to the owner. Additionally, more time was spent in areas close to the dispenser compared to the CST. This trend became more evident in the second pairing exposure (B2), where, in addition, a shortening of the routes was observed compared to the first pairing exposure (B1).

On the other hand, during extinction (A1 and A2), a different movement pattern was observed in all dogs. In general, restricted movement patterns were observed in areas close to the owner, as well as a smaller number of regions in which the accumulated time per session was distributed. This trend was more evident in the second extinction exposure (A2), and a restriction in movement patterns was also observed compared to the first extinction exposure (A1).

Figure 3 shows the total distance traveled per session for each dog. The X-axis indicates the experimental sessions and the Y-axis indicates the total distance traveled in centimeters. These graphs show that the mean values for distance traveled—the dotted line parallel to X—tended to be higher in pairing (B1 and B2) than in extinction (A1 and A2) for all three subjects. A progressive reduction in distance traveled was observed as the experiment progressed, especially in A2. The consistency between the mean values, level changes, tendencies, and variability per condition in most subjects (D1 and D2) suggests a robust effect of the type of contingencies in force (pairing or extinction) on the total distance traveled. As a complementary analysis, Figure 7, left panel, shows the values of distance traveled for each session for each of the subjects, which corroborate higher values during pairing compared to extinction. Quantitative analysis using LMM revealed a significant difference by condition (pairing/extinction) in this variable 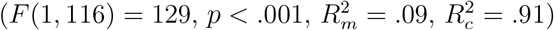

**Figure 3.**
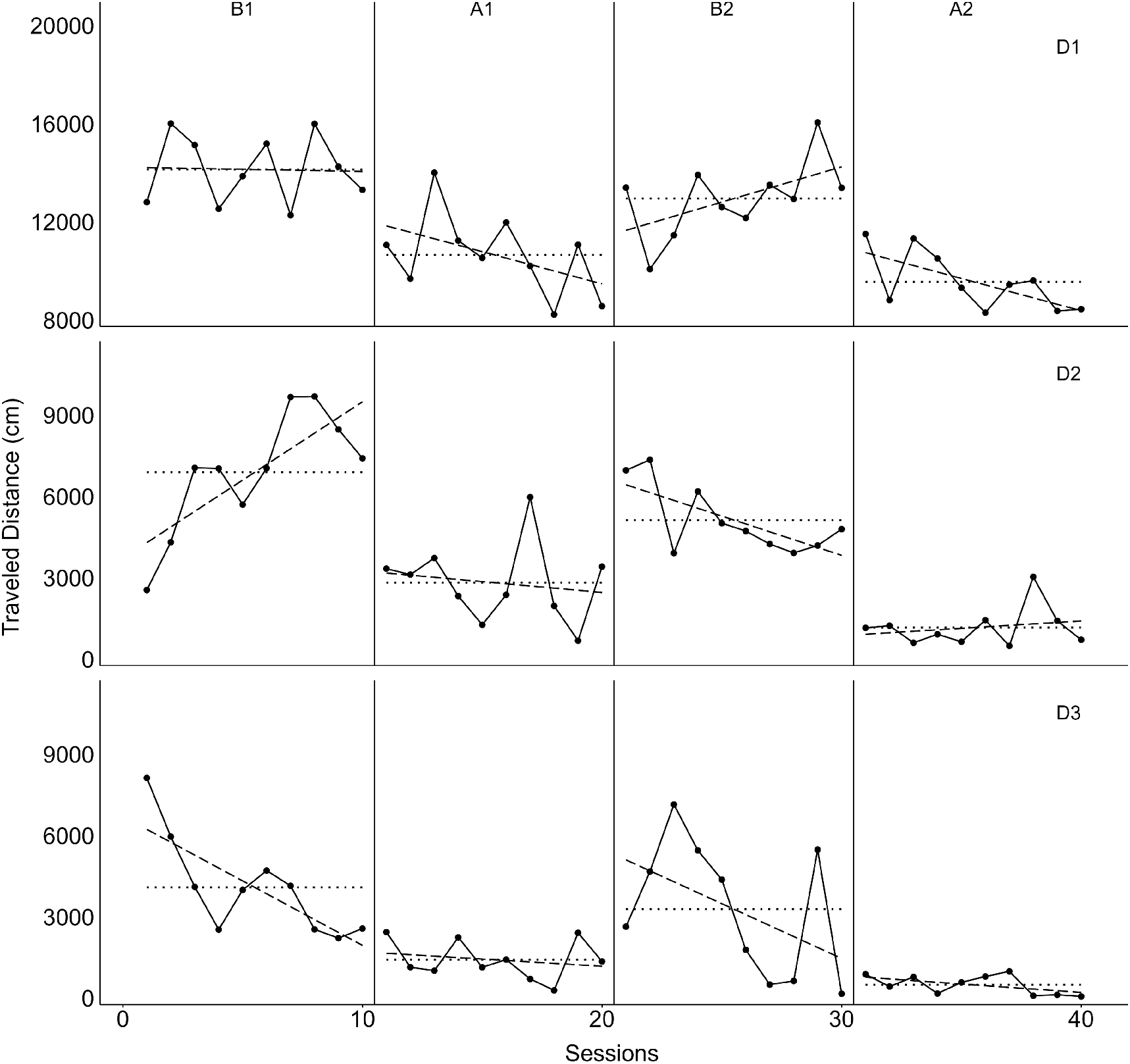
Total distance traveled per session. Each panel shows the total distance traveled (Y axis, 0-10,000 cm) throughout the sessions (X-axis) for each subject. The top, middle, and bottom panels correspond to D1, D2, and D3, respectively. The divisions within each panel represent the experimental conditions—Pairing (B) and Extinction (A)—ordered from left to right according to their sequence of exposure. The vertical lines indicate condition changes, while the horizontal dotted lines represent the mean values per condition. The dashed line shows the simple linear regression adjusted for each condition. Because D1 traveled a considerably greater distance than D2 and D3, the Y-axis of their panel was delimited independently (8000-20000 cm) to facilitate visual comparison with the other subjects.

Figure 4 shows the location entropy of the three subjects during the experimental sessions. The X-axis shows the experimental sessions and the Y-axis shows location entropy. These graphs show that the mean location entropy values tended to be higher in pairing (B1 and B2) than in extinction (A1 and A2) for all three subjects. This suggests that in the condition in which the tone-food contingency was in effect, the organisms tended to distribute their location among a greater number of regions within each session, while when the contingency relationship changed and only the tone (CS) was presented, the organism’s location tended to be more concentrated in a few regions. As a complementary analysis, Figure 7, right panel, shows the location entropy values for each session for each of the subjects, which corroborate higher values during pairing compared to extinction. Quantitative analysis using LMM revealed a significant difference in condition (pairing/extinction) in this variable 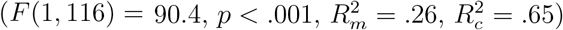.

**Figure 4.**
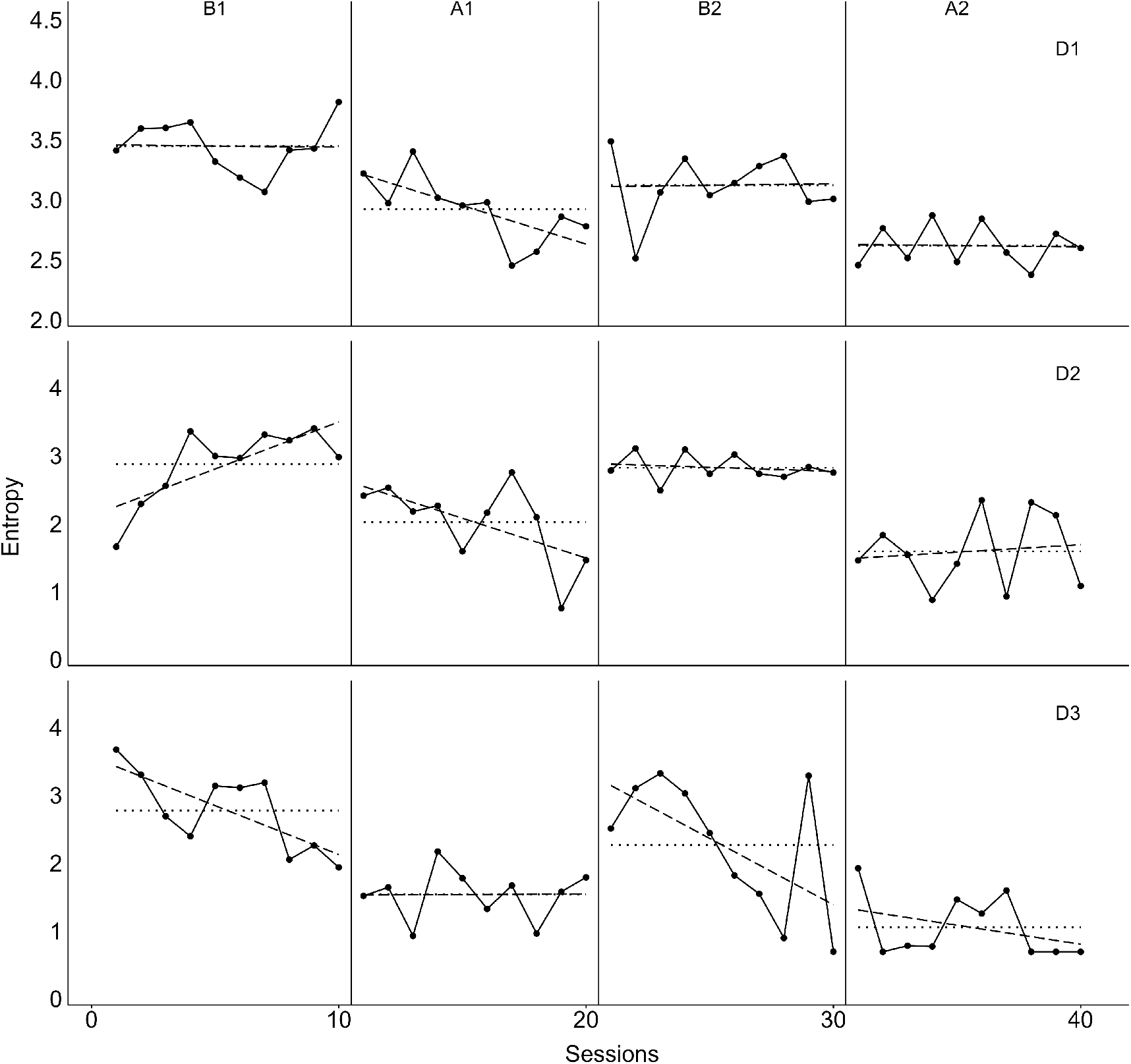
Location entropy. Each panel shows location entropy (Y axis, 0–4) across sessions (X axis) for each subject. The top, middle, and bottom panels correspond to D1, D2, and D3, respectively. The divisions within each panel represent the experimental conditions—Pairing (B) and Extinction (A)—ordered from left to right according to their sequence of exposure. The vertical lines indicate condition changes, while the horizontal dotted lines represent the mean values per exposure. The dashed line shows the simple linear regression adjusted for each condition. Because the location entropy values for D1 remained high, the Y-axis of its panel was delimited independently (2–4) to facilitate visual comparison with the other subjects.

Figure 5 shows the average distance (Y axis) to relevant objects (dispenser–owner) per session (X axis) for each subject. The brown series represent the mean distance values relative to the dispenser, while the green series represent the mean distance values relative to the owner. These graphs show a consistent pattern in the spatial organization of behavior based on programmed contingencies. In general, during the pairing (B1, B2), the distance to the dispenser was shorter, while proximity to the owner was reduced. In contrast, during extinction (A1, A2), the dogs increased their distance from the dispenser and increased their proximity to the owner. This effect suggests that the dogs’ location varied systematically depending on the type of contingencies, reflecting the general adjustment of their spatial behavior to the experimental conditions. In pairing, the functionally relevant areas were those close to the dispenser, established by contingency arrangement (tone–food), while in extinction, the areas close to the owner gained predominance. As a complementary analysis, Figure 8 shows the mean distance values to the dispenser and owner for each session for each of the subjects, which corroborate lower mean distance values to the dispenser during pairing compared to extinction. On the other hand, during extinction, lower mean distance values to the owner are corroborated in comparison with pairing. Quantitative analyses using LMM revealed a significant effect of condition on both mean distance to the dispenser 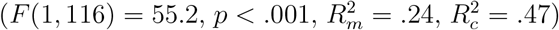, and mean distance to the owner 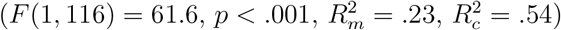.

**Figure 5.**
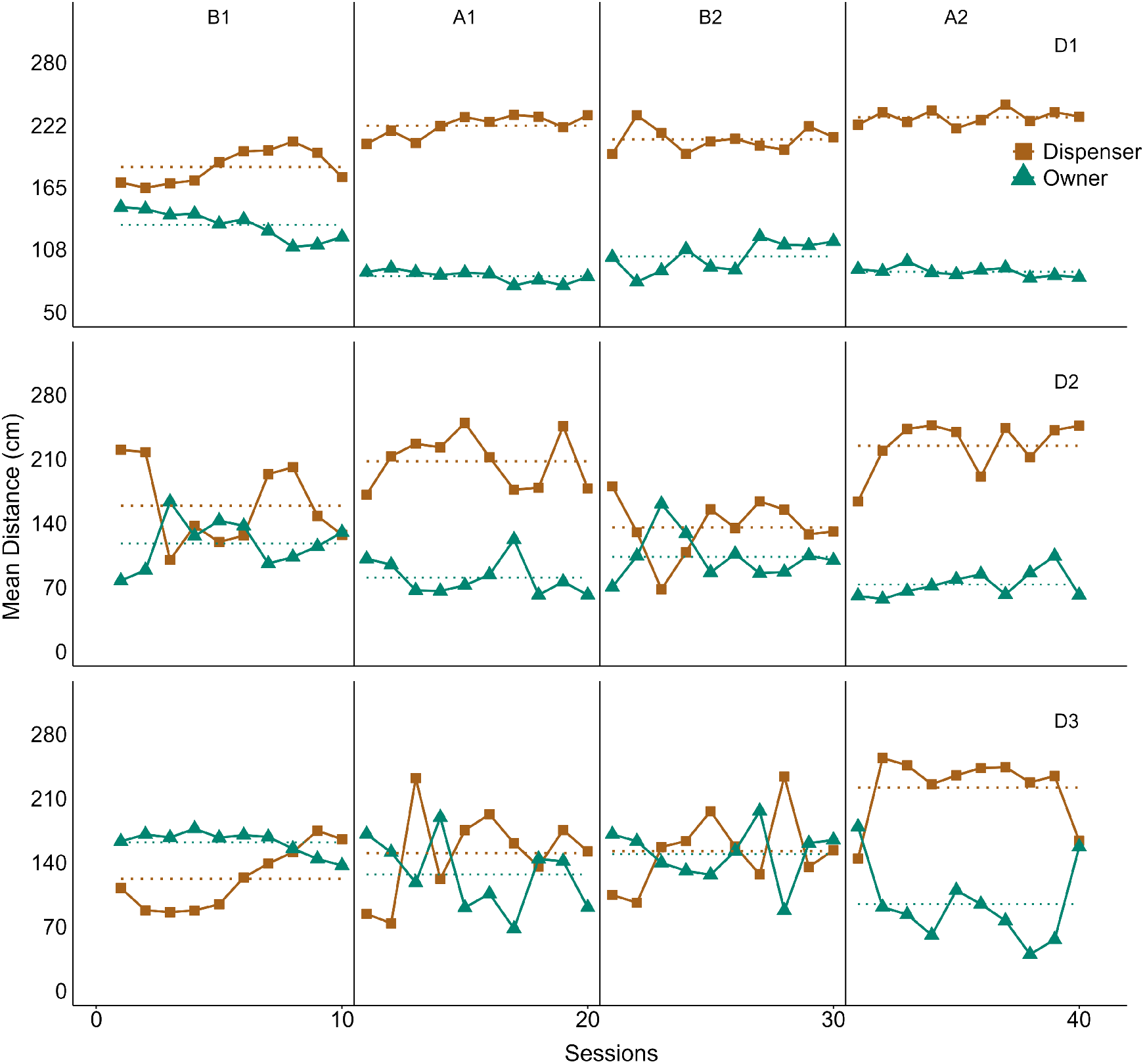
Mean distance to dispenser-owner. Each panel shows the mean distance to the dispenser-owner (Y-axis, 0–280 cm) across sessions (X-axis) for each subject. The top, middle, and bottom panels correspond to D1, D2, and D3, respectively. The divisions within each panel represent the experimental conditions—Pairing (B) and Extinction (A)—ordered from left to right according to their sequence of exposure. The solid brown line with filled squares indicates the mean distance to the dispenser per session, while the solid green line with filled triangles indicates the mean distance to the owner. The vertical lines indicate condition changes, while the horizontal dotted lines represent the mean values of distance to the dispenser (brown) and owner (green) per exposure. Because the D1 values were considerably high, the Y-axis of its panel was delimited independently (50–280 cm) to facilitate visual comparison with the other subjects.

Figure 6 shows the conditional approach index to the dispenser, which was calculated per trial and grouped by experimental condition (pairing/extinction). The Y-axis shows the index value, which ranges from 0 to 1. Each individual dot corresponds to a trial, color-coded according to the subject. The violin plots illustrate the distribution of data by condition.

**Figure 6.**
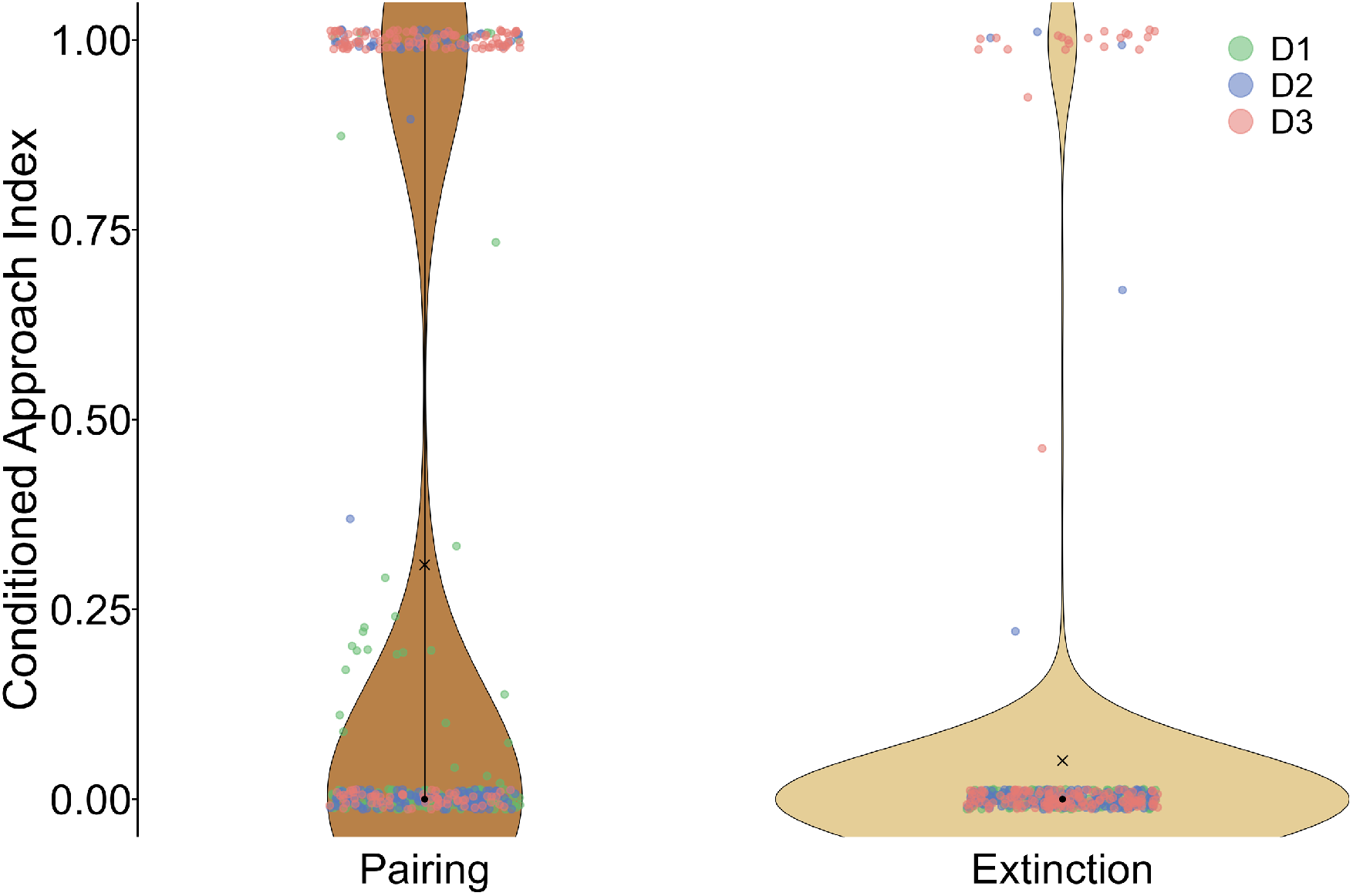
Conditional approach index to the dispenser by condition. Violin plots corresponding to the conditional approach index to the dispenser per trial. The violin with the highest saturation represents the pairing condition, while the clear one corresponds to the extinction condition. The black dot within each violin indicates the median, while the “X” represents the mean. The vertical line within each violin indicates the central 50% of the data (interquartile range, IQR). The green, blue, and red dots correspond to the individual data per trial of D1, D2, and D3, respectively. Statistical analysis (Mixed Linear Model considering the subject as a random effect) revealed a significant effect of condition 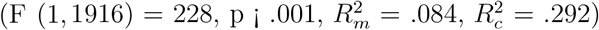. The data correspond to the trials of the last five sessions (6–10) of each exposure (N=480 per condition).

**Figure 7.**
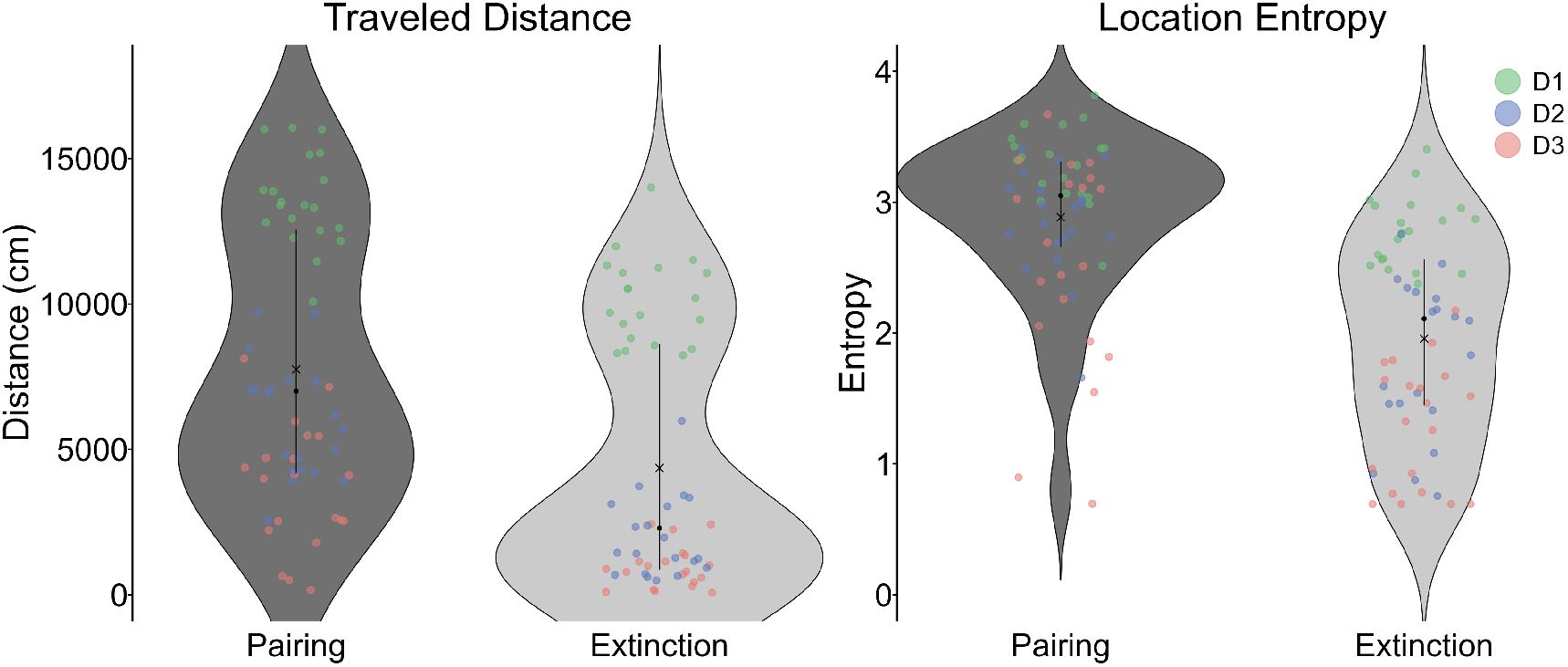
Comparison of total distance traveled and location entropy by condition. Violin plots corresponding to the total distance traveled per session (left panel) and location entropy per session (right panel), grouped by condition. The violins with greater saturation represent the pairing condition, while the lighter ones correspond to the extinction condition. The black dot within each violin indicates the median, while the “X” represents the mean. The vertical line within each violin indicates the central 50% of the data (interquartile range, IQR). The green, blue, and red dots in each graph correspond to the individual data for D1, D2, and D3, respectively. Statistical analysis (Mixed Linear Model considering the subject as a random effect) revealed a significant effect of condition for both total distance traveled 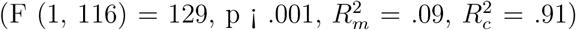 and location entropy 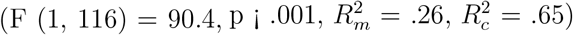. The data correspond to the total number of sessions of each subject throughout the experiment (N=60 per condition).

**Figure 8.**
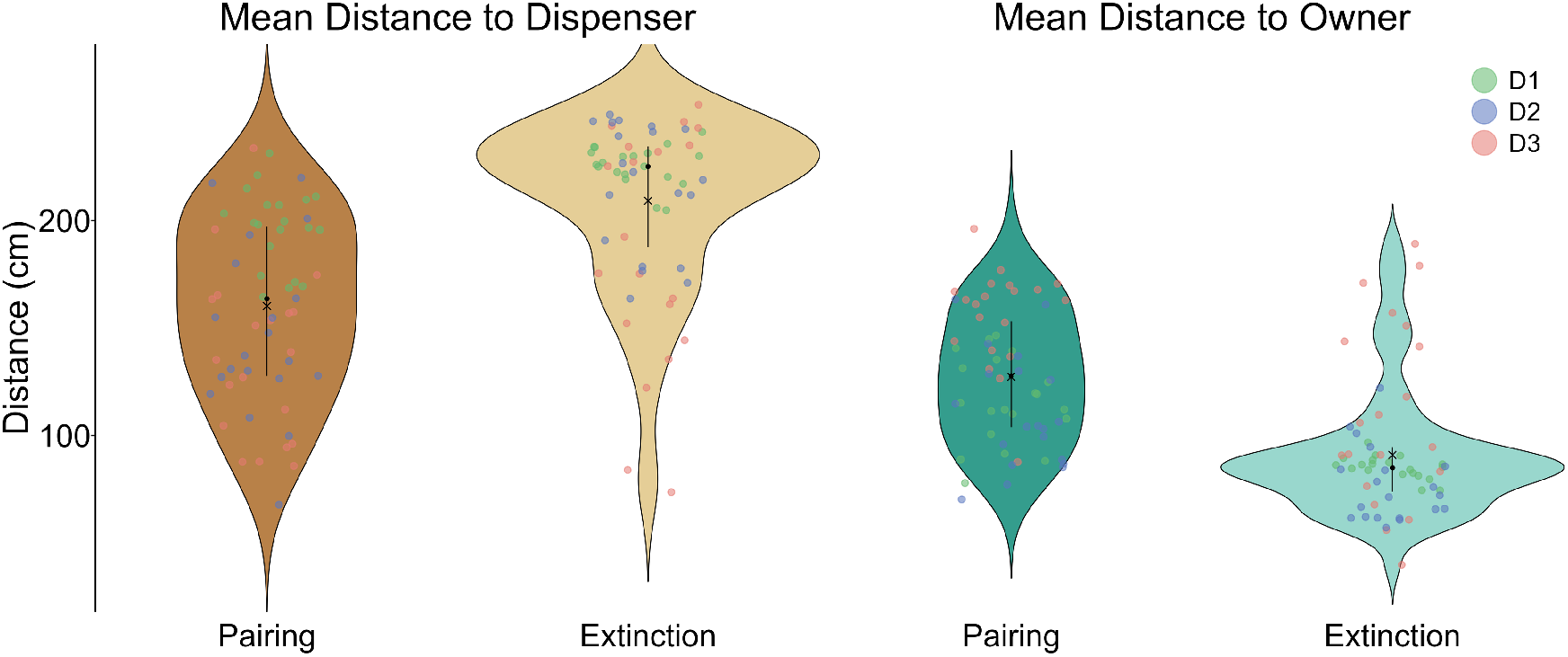
Comparison of mean distance to dispenser-owner by condition. Violin plots corresponding to the mean distance to the dispenser (brown violins) and owner (green violins), grouped by condition. The violins with greater saturation represent the pairing condition, while the lighter ones correspond to the extinction condition. The black dot within each violin indicates the median, while the “X” represents the mean. The vertical line within each violin indicates the central 50% of the data (interquartile range, IQR). The green, blue, and red dots in each graph correspond to the individual data for D1, D2, and D3, respectively. Statistical analysis (Mixed Linear Model considering the subject as a random effect) revealed a significant effect of condition for both mean distance to the dispenser 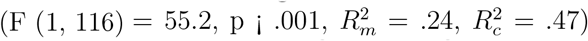 and for mean distance to the owner 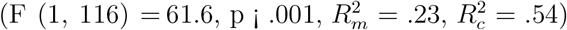. The data correspond to the total number of sessions for each subject throughout the experiment (N=60 per condition).

The results show a clear difference in experimental conditions. During the pairing condition, most trials resulted in extreme index values (close to 0 and 1), suggesting a marked alternation between trials with and without a conditional approach to the dispenser. In contrast, during the extinction condition, the values were predominantly concentrated around 0, indicating a systematic absence of a conditional approach to the dispenser. Quantitative analysis using LMM revealed a significant difference by condition on the conditional approach index 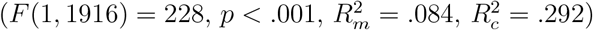.

## 4 Discussion

The objective of this study was to analyze changes in the spatiotemporal dynamics of the behavior of free-moving domestic dogs under appetitive Pavlovian contingencies and their extinction.

In pairing, the subjects tended to remain and move in areas close to the dispenser, alternating with areas close to the owner or on the periphery of the experimental room. They also traveled a greater distance and moved more extensively within each session. In addition, the consistent establishment of a conditional approach pattern to the dispenser was observed. In contrast, during extinction, the subjects moved around areas close to the owner or on the periphery, covered less distance, and showed less variability in locomotion. In addition, the pattern of conditional approach to the dispenser was reduced.

A key finding was that, during pairing, the regions near the dispenser directed the dogs’ behavior, which was evident from the repetition of the location and the narrowing of activity patterns in those regions. In contrast, when food delivery ceased (extinction) and the contingent relationship (place-tone-food) was interrupted, a reorganization of behavior and eventual disappearance of the patterns established in pairing was observed. These findings are plausible indications of the CS function acquired by the areas near the dispenser in pairing and are consistent with the findings reported by Kupalov (Kupalov, 1969, 1983) in his research on conditional place or situational reflexes.

It should be noted that, in the present study, unlike Kupalov’s work, the use of a fixed-time program for presenting stimuli (independent of the organism’s activity pattern) made it possible to methodologically ensure the organization of Pavlovian contingencies, marginalizing operant processes or interactions between different types of contingencies (operant and Pavlovian).

In addition to the central effects attributed to the CS-US contingency, other displacement patterns were identified, which, although not the main focus of the study, contribute to characterizing the spatiotemporal dynamics of the subjects’ behavior. During pairing, additional displacement patterns were identified, centered on regions close to the owner or peripheral regions, particularly in a corner corresponding to the door of the experimental room. In contrast, during extinction, movement patterns were concentrated specifically in regions close to the owner and the door. These findings suggest that the spatiotemporal dynamics of the observed behavior were not determined exclusively by the regions associated with the CS-US contingency, but were the result of multiple spatial relationships present in the experimental context.

It is reasonable to argue that, during the pairing condition, not only did the areas near the dispenser influence the direction of the subjects’ movement, but also those functionally relevant regions derived both from the subject’s behavioral history—particularly their interaction with the owner—and from ecological-behavioral patterns specific to domestic dogs. These include dog-human interspecific proximity and the exploration of areas of potential relevance, such as the enclosure door. The convergence of multiple functional regions was reflected in movement patterns characterized by back-and-forth trajectories, as well as alternating stays between these areas, which led to higher values of location entropy and distance traveled. In contrast, during the extinction condition, the areas near the owner’s location and the door became more functionally relevant, being the main determinants of the observed spatiotemporal dynamics.

The plausibility of this interpretation is reinforced by previous literature on the spatial behavior of domestic dogs. Previous studies (Völter et al., 2023) have shown that, even in arrangements with multiple relevant zones (e.g., toys, owners, and strangers to the dogs), the areas close to humans and the door of the experimental enclosure maintain a high frequency of visits. Added to this is the species’ ability to orient itself based on spatial markers or landmarks (Milgram et al., 2002; Muscosky and Horowitz, 2024). It is also relevant to consider that, in domestic contexts, where humans organize the dogs’ environment in relation to the availability of food (US) (Udell and Wynne, 2008), the presence or location of the owners could function as CS, directing the dogs’ spatial behavior.

The spatial organization of behavior observed in the present study falls within a broader framework of empirical evidence. Previous research has documented similar patterns in other species, such as rats, in which low contiguity between functionally relevant regions—for example, areas near the feeder and areas that serve as potential refuges, including corners— promotes alternating movement among these areas (León et al., 2020).

Furthermore, the results of this study coincide with evidence of the appearance of conditional approach–withdrawal patterns in other non-human animals such as pigeons and rodents (Fitzpatrick et al., 2013; Hearst et al., 1980; Wasserman et al., 1974, among others). In the case of the present study, the conditional approach index used and applied in different studies in the field (Hearst et al., 1980; Wasserman et al., 1974), allowed us to differentiate clear changes in the pattern of approach to the dispenser during the presentation of the CS, depending on the type of condition.

Taken together, the findings suggest that the spatial dimension of behavior is a relevant variable that participates in basic learning processes in different species and becomes important from a comparative perspective (Balcı et al., 2025; Hernández-Eslava et al., 2023).

Although the main contribution of this study is empirical and methodological, the findings— in particular, greater distance traveled and location entropy in pairing—allow us to propose possible extensions to the field of animal welfare. Excessive confinement, lack of exercise, and low environmental stimulation have been documented as common sources of stress and compulsive behaviors in domestic dogs (Lindsay, 2001). In this sense, replicating the arrangement used in this study could function as a form of environmental enrichment (Hunt et al., 2022) to increase physical activity and behavioral variability. This would be consistent with evidence showing the effectiveness of non-contingent reinforcement programs in reducing undesirable behaviors in shelter dogs (Protopopova and Wynne, 2015).

Although the findings of this study are robust and the possible applications plausible, it is important to consider some limitations, which are described below.

First, although the conditional approach index used (Hearst et al., 1980; Wasserman et al., 1974) revealed a certain synchrony between the subjects’ movements and the presentation of the CS, it did not allow for the precise identification of behavioral changes before and after each trial. Therefore, new quantification strategies are needed to distinguish more finely between behavioral patterns caused by the presentation of the CS (conditional response), those derived from the passage of time (e.g., temporal conditioning), and those derived from the presentation of the US (unconditional response).

Furthermore, a progressive decrease in D3’s engagement with the task was observed throughout the experiment. This effect could have been due both to the low value of the food used as US—probably due to its low palatability for the subject—and to the number of sessions conducted, which may have been high for D3.

Finally, the behavioral recording in this study was based on a two-dimensional tracking system with a single reference point (center of mass), which prevented the recording of additional behaviors, such as orientation responses (Sokolov, 1982) or postures. Previous research (Barnard et al., 2016; Völter et al., 2023) has shown the usefulness of multipoint tracking and three-dimensional video recording for identifying a wide range of behaviors in domestic dogs. Incorporating these technologies into future studies could enable a more accurate and comprehensive characterization of the spatiotemporal organization of organisms’ behavior across different contingency arrangements (Balcı et al., 2025).

In conclusion, this study provides novel evidence on the spatiotemporal dynamics of domestic dog behavior under Pavlovian contingencies and highlights the usefulness of tracking tools in behavioral analysis. Beyond the effects observed in a particular species, the findings open new avenues for empirically exploring how space is functionally integrated into contingent relationships that incorporate temporal arrangements, giving rise to the establishment of distinctive spatial behavior patterns.

## Acknowledgements

The authors would like to thank Fryda Díaz for her valuable support in programming the experimental event control software for this study.

The first author was supported by a graduate scholarship (No. 800045) from the Secretaría de Ciencia, Humanidades, Tecnología e Innovación (Móxico).

## Notes

### Competing Interest Statement

The authors have declared no competing interest.

## References

Balcı, F., Hernandez, V., Hoşer, A., and Leén, A. (2025). Movement path as an ethological lens into interval timing. Behavioural Processes, 230:105233.

Barnard, S., Calderara, S., Pistocchi, S., Cucchiara, R., Podaliri-Vulpiani, M., Messori, S., and Ferri, N. (2016). Quick, accurate, smart: 3d computer vision technology helps assessing confined animals’ behaviour. PloS one, 11(7):e0158748.

Bleuer-Elsner, S., Zamansky, A., Fux, A., Kaplun, D., Romanov, S., Sinitca, A., Masson, S., and van der Linden, D. (2019). Computational analysis of movement patterns of dogs with adhd-like behavior. Animals, 9(12):1140.

Brown, P. L. and Jenkins, H. M. (1968). Auto-shaping of the pigeon’s key-peck. Journal of the Experimental Analysis of Behavior, 11(1):1–8.

Buzsaki, G. (1982). The “where is it?” reflex: autoshaping the orienting response. Journal of the Experimental Analysis of Behavior, 37(3):461–484.

Carcassi, G., Aidala, C. A., and Barbour, J. (2021). Variability as a better characterization of shannon entropy. European Journal of Physics, 42(4):045102.

Domjan, M. (2010). Principios de aprendizaje y conducta. Wadsworth Cengage Learning, 6 edition.

Fernandez, E. J., Anderson, W., and Kowalski, A. (2023). Evaluation of an automated response-independent schedule on the behavioral welfare of shelter dogs. Journal of the Experimental Analysis of Behavior, 120(1):50–61.

Feuerbacher, E. N. and Wynne, C. D. L. (2011). A history of dogs as subjects in north american experimental psychological research. Comparative Cognition Behavior Reviews, 6:46–71.

Fitzpatrick, C. J., Gopalakrishnan, S., Cogan, E. S., Yager, L. M., Meyer, P. J., Lovic, V., Saunders, B. T., Parker, C. C., Gonzales, N. M., Aryee, E., Flagel, S. B., Palmer, A. A., Robinson, T. E., and Morrow, J. D. (2013). Variation in the form of pavlovian conditioned approach behavior among outbred male sprague-dawley rats from different vendors and colonies: sign-tracking vs. goal-tracking. PloS one, 8(10):e75042.

Hearst, E. (1975). Pavlovian conditioning and directed movements. Psychology of Learning and Motivation, 9:215–262.

Hearst, E., Bottjer, S. W., and Walker, E. (1980). Conditioned approach-withdrawal behavior and some signal-food relations in pigeons. Bulletin of the Psychonomic Society, 16(3):183– 186.

Henton, W. W. and Iversen, I. H. (1978). Classical Conditioning and Operant Conditioning: A Response Pattern Analysis. Springer-Verlag.

Hernández-Eslava, V., Leén, A., Guzmán, I., DÍaz, F., Avendaño-Garrido, M. L., Toledo, P., et al. (2023). An ecological approach to the effects of water-source locations and time-based schedules on entropy and spatio-temporal behavioral features. International Journal of Comparative Psychology, 36.

Howse, M. S., Anderson, R. E., and Walsh, C. J. (2018). Social behaviour of domestic dogs (canis familiaris) in a public off-leash dog park. Behavioural Processes, 157:691–701.

Hunt, R. L., Whiteside, H., and Prankel, S. (2022). Effects of environmental enrichment on dog behaviour: Pilot study. Animals, 12(2):141.

Jenkins, H. M., Barrera, F. J., Ireland, C., and Woodside, B. (1978). Signal-centered action patterns of dogs in appetitive classical conditioning. Learning and Motivation, 9(3):272– 296.

Kolosova, T. E., Fomicheva, E. E., and Obukhova, G. P. (1982). Situational conditioned reflexes in intact and callosectomized dogs: Effects of changes in location of conditioned and unconditioned stimuli. Neuroscience and Behavioral Physiology, 12(6):457–462.

Kupalov, P. S. (1969). The formation of conditioned place reflexes. In Cole, M. and Maltzman, I., editors, A Handbook of Contemporary Soviet Psychology, pages 735–762. Basic Books.

Kupalov, P. S. (1983). Situational conditional reflexes: Physiologic studies of the higher nervous activity of freely moving animals. Pavlovian Journal of Biological Science, 18(1):13– 21.

Kupalov, P. S., Voevodina, O. N., Volkova, V. D., Malyukova, I. V., and Selivanova, A. T. (1964). Situational Conditioned Reflexes in Dogs in Normal and Pathological State. Clearinghouse.

Lanovaz, M. J., Hernandez, V., and Leén, A. (2025). Machine learning to detect schedules using spatiotemporal data of behavior: A proof of concept. Journal of the Experimental Analysis of Behavior, 124(1):e70029.

Leén, A., Hernandez, V., Lopez, J., Guzman, I., Quintero, V., Toledo, P., Avendaño-Garrido, M. L., Hernandez-Linares, C. A., and Escamilla, E. (2021). Beyond single discrete responses: An integrative and multidimensional analysis of behavioral dynamics assisted by machine learning. Frontiers in Behavioral Neuroscience, 15:681771.

Leén, A., Hernández, V., Huerta, U., Hernández-Linares, C. A., Toledo, P., Avendaño Garrido, M. L., Escamilla Navarro, E., and Guzmán, I. (2020). Ecological location of a water source and spatial dynamics of behavior under temporally scheduled water deliveries in a modified open-field system: An integrative approach. Frontiers in Psychology, 11:577903.

Lindsay, S. R. (2000). Handbook of applied dog behavior and training: Adaptation and learning. Blackwell Publishing.

Lindsay, S. R. (2001). Handbook of Applied Dog Behavior and Training: Etiology and Assessment of Behavior Problems. Blackwell Publishing.

Mehrkam, L. R., Perez, B. C., Self, V. N., Vollmer, T. R., and Dorey, N. R. (2020). Functional analysis and operant treatment of food guarding in a pet dog. Journal of Applied Behavior Analysis, 53(4):2139–2150.

Milgram, N. W., Head, E., Muggenburg, B., Holowachuk, D., Murphey, H., Estrada, J., Ikeda-Douglas, C. J., Zicker, S. C., and Cotman, C. W. (2002). Landmark discrimination learning in the dog: effects of age, an antioxidant fortified food, and cognitive strategy. Neuroscience and Biobehavioral Reviews, 26(6):679–695.

Muscosky, L. and Horowitz, A. (2024). Distinguishing doors and floors on all fours: Landmarks as tools for vertical navigation learning in domestic dogs (canis familiaris). Animals, 14(22):3316.

Pavlov, I. P. (1927). Conditioned Reflexes: An Investigation of the Physiological Activity of the Cerebral Cortex. Oxford University Press.

Pear, J. J. (1985). Spatiotemporal patterns of behavior produced by variable-interval schedules of reinforcement. Journal of the Experimental Analysis of Behavior, 44(2):217–231.

Protopopova, A. and Wynne, C. D. L. (2015). Improving in-kennel presentation of shelter dogs through response-dependent and response-independent treat delivery. Journal of Applied Behavior Analysis, 48(3):590–601.

Schielzeth, H., Dingemanse, N. J., Nakagawa, S., Westneat, D. F., Allegue, H., Teplitsky, C., Réale, D., Dochtermann, N. A., Garamszegi, L. Z., and Araya-Ajoy, Y. G. (2020). Robustness of linear mixed-effects models to violations of distributional assumptions. Methods in Ecology and Evolution, 11(9):1141–1152.

Sokolov, Y. N. (1982). Percepcién y reflejo condicionado. Trillas. Original work published 1963.

Stepień, I. (1974). The magnet reaction, a symptom of prefrontal ablation. Acta Neurobiologiae Experimentalis, 34(1):145–160.

Tarpy, R. M. (2000). Aprendizaje: TeorÍa e investigacién contemporáneas. McGraw-Hill. Original work published 1967.

Udell, M. A. R. and Wynne, C. D. L. (2008). A review of domestic dogs’ human-like behaviors: or why behavior analysts should stop worrying and love their dogs. Journal of the Experimental Analysis of Behavior, 89(2):247–261.

Völter, C. J., Starić, D., and Huber, L. (2023). Using machine learning to track dogs’ exploratory behaviour in the presence and absence of their caregiver. Animal Behaviour, 197:97–111.

Wasserman, E. A., Franklin, S. R., and Hearst, E. (1974). Pavlovian appetitive contingencies and approach versus withdrawal to conditioned stimuli in pigeons. Journal of Comparative and Physiological Psychology, 86(4):616–627.

Zamansky, A., Bleuer-Elsner, S., Masson, S., Amir, S., Magen, O., and van der Linden, D. (2018). Effects of anxiety on canine movement in dog-robot interactions. Animal Behavior and Cognition, 5(4):380–387.

Zener, K. (1937). The significance of behavior accompanying conditioned salivary secretion for theories of the conditioned response. American Journal of Psychology, 50:384–403.

